# Mast cell-mediated inflammation relies on insulin-regulated aminopeptidase controlling cytokine export from the Golgi

**DOI:** 10.1101/2022.01.21.477149

**Authors:** Mirjana Weimershaus, Caroline Carvalho, Rachel Rignault, Emmanuelle Waeckel-Enee, Michael Dussiot, Peter van Endert, Thiago Trovati Maciel, Olivier Hermine

## Abstract

Upon activation, mast cells rapidly release preformed inflammatory mediators from large cytoplasmic granules via regulated exocytosis. This acute degranulation is followed by a late activation phase involving synthesis and secretion of cytokines, growth factors and other inflammatory molecules via the constitutive pathway that remains ill-defined. Here we describe a role for an insulin-responsive vesicle-like endosomal compartment, marked by insulin-regulated aminopeptidase (IRAP), in the secretion of TNF-α- and IL-6 in mast cells and macrophages. IRAP-deficient mice are protected from TNF-α-dependent kidney injury and inflammatory arthritis. In the absence of IRAP, TNF-α fails to be efficiently exported from the Golgi. Subsequently, reduced co-localization of VAMP3+ endosomes with Stx4 was observed, while VAMP8-dependent exocytosis of secretory granules was facilitated. Chemical targeting of IRAP+ endosomes reduced pro-inflammatory cytokine secretion thereby highlighting this compartment as a promising target for the therapeutic control of inflammation.

## Introduction

Mast cells are phylogenetically ancient innate immune cells residing in most connective and mucosal tissues. Their prominent characteristic are large cytoplasmic granules filled with proteases, histamine, serotonin, cytokines and inflammatory mediators that are rapidly released upon signaling through specific cell surface receptors, including FcεR, complement receptors, TLRs, and GPCRs. Moreover, in a second phase after activation, mast cells synthesize and secrete large amounts of pro- and anti-inflammatory cytokines, chemokines and growth factors, prostaglandins and leukotrienes. Interestingly, certain stimuli, such as LPS and IL-1β exclusively activate secretion of *de novo* synthesized mediators without triggering release of preformed granules ^1–3^. While our understanding of the processes involved in the biogenesis and triggered release of preformed granule content in anaphylactic reactions has been advanced in recent years, much remains to be learned about the immunomodulatory roles of mast cell-derived cytokines, chemokines and growth factors, including the regulation of their secretion ^3,4^.

Mast cell-derived cytokines, chemokines and growth factors can act in autocrine, paracrine, local and systemic fashion, and are involved in physiological and protective processes such as angiogenesis, wound healing and the immune defense against bacteria and viruses, as well as in pathological processes such as autoimmune, metabolic and neurological disorders, fibrosis and cancer ^5^. With regards to the role of mast cells in disease-related settings, pro-inflammatory cytokines play a prime role. Especially, mast cell-derived TNF-α and IL-6 and have been in the focus of numerous studies. They have been reported to chemotactically attract neutrophils and macrophages, to upregulate adhesion molecules in endothelial cells ^6–8^, modulate DC functions ^9,10^, promote colitis ^11^, mediate cisplatin-induced kidney injury ^12^, participate in inflammatory arthritis ^13^, and control airway hyperreactivity, inflammation, and Th2 recruitment and cytokine production in a murine antigen-induced asthma model ^14^. Targeting cytokine synthesis and secretion in the late or chronic activation phase in mast cells might therefore represent a promising therapeutic strategy to prevent adverse mast cell-related inflammatory reactions and actively shape the immuno-modulatory responses of these versatile cells.

Of note, several hundreds of biological compounds with various functions have been identified in the mast cell secretome ^15,16^. Some of these have opposed physiological functions, suggesting that secretory pathways and products in mast cells may be temporally and spatially regulated.

Exocytic mechanisms, *i*.*e*. active vesicular transport resulting in compound release, are present in all eukaryotic cells. While products and biological functions of exocytosis vary largely between cell types, the underlying pathways and trafficking machinery are highly conserved. Two major pathways can be mechanistically distinguished, referred to as regulated and constitutive secretion, respectively. A few recent studies have started to elucidate the mechanistic aspects that segregate constitutive cytokine secretion from regulated granule exocytosis ^17–20^. This discrimination has been advanced by the identification of the distinct Soluble N-ethylmaleimide-sensitive factor attachment protein receptors (SNARE) proteins involved in either pathway. SNARE proteins confer membrane identity and control fusion of lipid bilayers of distinct compartments, as each v (vesicular)-SNARE family member can form complexes only with a limited set of t (target)-SNARE partners.

In the constitutive pathway in macrophages, where cytokine secretion has been studied best, following translation, cytokines traffic from the ER through the Golgi stacks, from where they are exported in syntaxin (Stx) 6 SNARE-identified vesicles ^21^. These post-Golgi carriers then fuse with Rab11+ VAMP3+ recycling endosomes and are further transported to the plasma membrane (PM) ^21,22^. The major SNARE complex required for VAMP3 vesicle fusion with the PM is composed of SNAP23 and Stx4 in murine mast cells and macrophages ^19,23^. As specific sorting signals routing cargo from the trans-Golgi network (TGN) into constitutive secretion are yet to be identified, this pathway is considered as a “default” pathway ^24^.

In contrast, proteins destined to packing into secretory lysosome-related granules are actively sorted away in the Golgi to form immature secretory granules. These pre-granules undergo a series of fusion and fission events which result in removal of mis-sorted cargo and condensation giving rise to mature granules stored in the cytoplasm ^25^. Release of secretory granules in mast cells requires activation via inflammatory cell surface receptors, including FcεR, FcγR, TLRs, complement receptors and other GPCRs, and is mediated through elevation of intracellular Ca^2+^ levels and activation of PKC ^26,27^. The main SNARE protein mediating attachment and fusion of the secretory granules with the plasma membrane is VAMP8. Of note, the same SNARE complex as in the constitutive pathway, composed of SNAP23 and Stx4, is employed for secretory granule attachment and fusion at the plasma membrane ^19,28,29^.

Interestingly, limited amounts of preformed cytokines including TNF-α are found in secretory granules ^6^ and, at least in human mast cell lines, have been suggested to traffic there by re-endocytosis from the extracellular space rather than after direct TGN-sorting to these granules ^30^.

Moreover, a distinct compartment of recycling vesicles has been described in mast cells ^31^. These vesicles are identified by the expression of insulin-regulated aminopeptidase (IRAP) and, in resting cells, exhibit a prevailing cytosolic distribution near the ER-Golgi intermediate compartment (ERGIC) from where they undergo slow recycling to the PM. Upon signal transduction through the FcεR, IRAP rapidly translocates to the plasma membrane, where it may participate in signal transduction events. Importantly, IRAP endosome mobilization is mechanistically segregated from the exocytosis of secretory granules ^31^. This suggests that IRAP and granule-contained mediators such as histamine are present in distinct compartments, but the precise relationship of IRAP to the different secretory pathways in mast cell is unknown. The possible positive or negative regulation of the degranulation process by IRAP endosomes, for instance, has not been elucidated by Liao et al. due to the absence of IRAP-invalidated cells or animals in the study. Thus, considering the intersection of exocytosis with endocytic and recycling pathways, as well as the importance of their coordination for mast cell physiology, we set out to investigate the communication of IRAP-containing endosomes with the aforementioned exocytic pathways.

IRAP endosomes have mainly been described as Glucose-transporter (Glut) 4 storage vesicles (GSV) in adipocytes and muscle cells where they have been extensively studied with respect to their function in insulin-stimulated Glut4 trafficking ^32,33^. In insulin-responsive cells, upon an activating signal through the insulin receptor, the dynamic retention of GSV in the cytosol is released, which allows them to traffic to the cell surface, thereby inserting Glut4, IRAP and other transmembrane GSV proteins into the plasma membrane ^34,35^. From here, IRAP and Glut4 are re-internalized into sorting endosomes, where they have been shown to interact with the retromer complex that promotes their deviation from the degradative late endosomal/lysosomal pathway and retrieves them for retrograde TGN ^36^, the GSV assembly and budding site.

Of note, in contrast to the relatively restricted Glut4 expression patterns to insulin-responsive tissues, IRAP-containing endosomes are widely expressed amongst cell types and tissues, where they are mobilized by cell-specific surface receptor signaling and employed for various cell type-specific functions ^37–39^. In this line, with regards to immune cells, the trafficking of IRAP endosomes has recently been recognized to intersect with and regulate phagosome maturation and MHC-I cross-presentation in dendritic cells ^40–43^, activation of TLR9 ^44^, as well as endo-and exocytic trafficking in T cells for the supply of TCR signaling components and optimal TCR signaling ^45^.

Here we report that IRAP endosomes are required for the post-Golgi transport of the pro-inflammatory cytokines TNF-α and IL-6 in the constitutive secretion pathway in mast cells *in vitro* and *in vivo* while limiting excessive degranulation of preformed mediators.

## Results

### IRAP endosomes are dispensable for secretory granule exocytosis

Aiming to shed light on the role of IRAP endosomes in mast cell exocytic trafficking pathways, we first colocalized IRAP with different endosomal markers of early and GSV-like endosomes (Fig. 1A). Like in dendritic cells, IRAP colocalized well with the early endosomal markers EEA1 and endocytized transferrin in mast cells, as well as with the GSV markers Rab14 and Stx6 involved in Golgi-to-endosome trafficking, confirming a high level of conservation of the IRAP-related vesicular trafficking machinery amongst different cell types. Interestingly, in activated cells, IRAP strongly colocalized with the granule-contained monoamine serotonin at the plasma membrane (Fig. 1B), similar to the observation with regards to histamine in the initial study ^31^, which prompted us to re-examine the role of IRAP in mast cell degranulation.

**Fig. 1.**
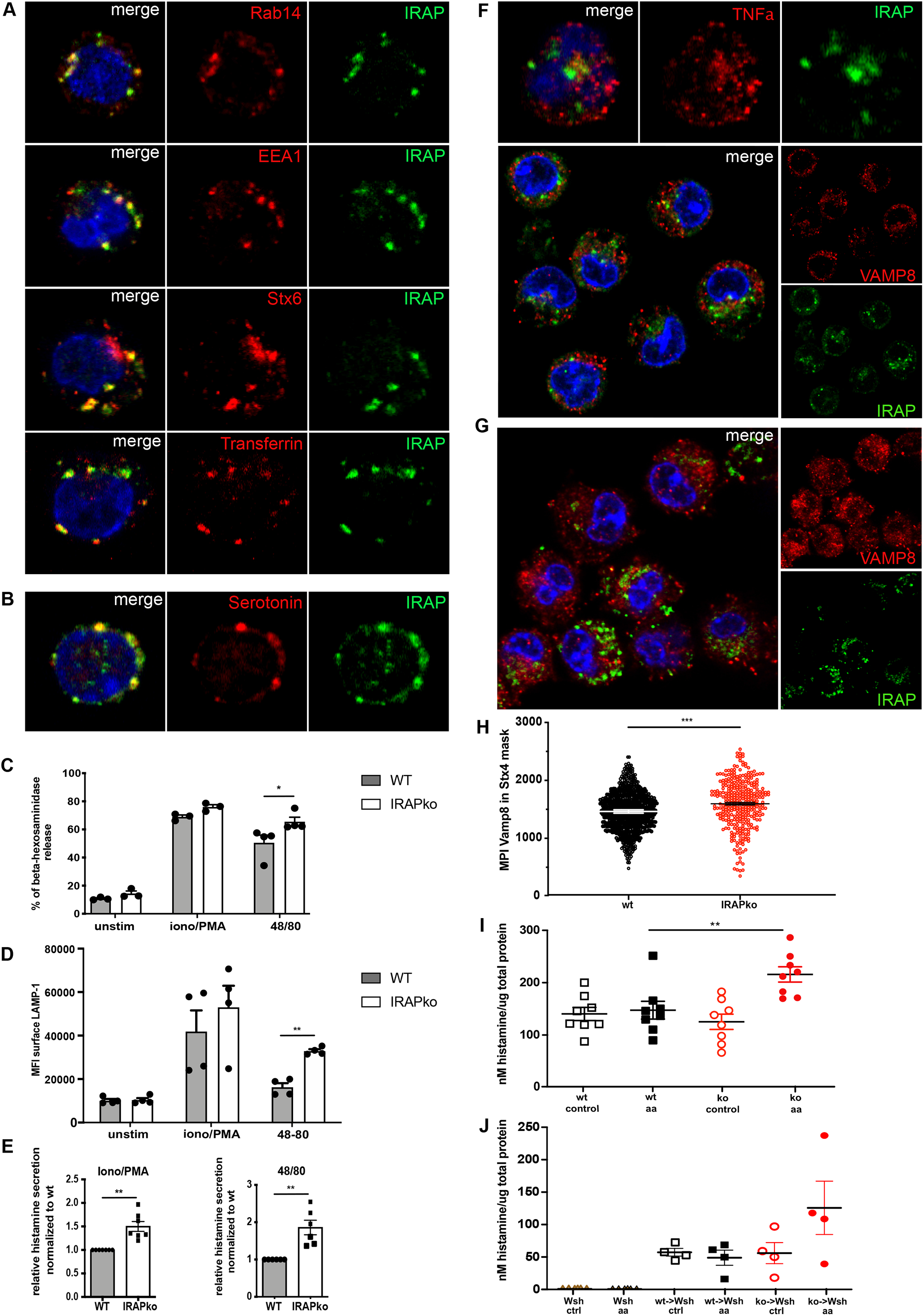
IRAP is dispensable for degranulation in mast cells. **(A)** Representative immunofluorescent images of resting BMMCs stained for IRAP and the endosomal markers Rab14, Stx6, EEA1, or 15min endocytized transferrin-AF647. **(B)** BMMC were stimulated for 30min with ionomycin/PMA prior to intracellular staining for serotonin and IRAP. **(C)** Beta-hexosaminidase secretion of peritoneal mast cells after 30min stimulation with ionomycin/PMA or 48/80 or without stimulation. Shown are mean values ± SEM of three independent experiments. **(D)** Degranulation analysis by flow cytometry via LAMP-1 surface staining of live peritoneal mast cells stimulated for 30min with ionomycin/PMA or 48/80 or left unstimulated. Shown are mean values ± SEM of three independent experiments. **(E)** Histamine secretion by peritoneal mast cells stimulated for 30min with ionomycin/PMA or 48/80. Shown are mean values ± SEM of the secretion of IRAPko cells normalized to wt from three independent experiments. **(F)** Resting BMMCs were fixed and immunostained for IRAP and VAMP8 or TNF-α. Images are representative of four independent experiments. **(G)** BMMCs were stimulated with ionomycin/PMA for 15min prior to fixation and immunostaining for VAMP8 and IRAP. Images are representative of two independent experiments. **(H)** Imaging flow cytometry quantification of VAMP8 mean pixel intensity (MPI) in the Stx4 mask of ionomycin/PMA-stimulated BMMC. Analysis was performed on >300 cells. **(I)** IRAPwt and ko mice or (J) mast-cell deficient Wsh mice reconstituted with IRAPwt and ko BMMC were challenged on one ear with 30mg/ml arachidonic acid while the control ear was left untreated. Histamine concentration was quantified in ear tissue homogenates and normalized to total protein concentration. Data are shown from one out of two experiments. *p<0.05,** p<0.01, ***p<0.001 for all figures.

Physiologically, ligation of mast cell surface receptors, including crosslinking of FcεR through IgE and cognate antigen, activates signaling cascades most of which converge to Ca^2+^ release from intracellular stores. If and how secretion of pre-stored granules versus *de novo* synthesized mediators upon Ca^2+^ signaling is regulated, is unknown.

As we aimed to analyze the potential involvement of IRAP endosomes in exocytosis separately from its hypothetical “upstream” role in the FcεR-related signaling cascade ^31^, we exclusively used FcεR-independent activation of mast cells throughout our study.

We stimulated peritoneal mast cells with ionomycin/PMA or with the GPCR-dependent compound 48/80 ^46^, and measured degranulation either as the release of the major granule component beta-hexosaminidase into the culture supernatant (Fig. 1C), or exocytosis of the lysosomal marker LAMP-1 in a flow cytometry assay (Fig. 1D). Degranulation was significantly increased in IRAP knock-out (IRAPko) cells upon 48/80 activation while the ionomycin/PMA stimulation led to strong degranulation responses without significant differences between IRAP-expressing and -deficient cells. As saturation effects may hide differences upon stimulation by ionomycin/PMA, we turned to an *in vitro* assay for the measurement of histamine release. This FRET-based technique is quantitative over a large range of histamine concentrations and detected significantly increased degranulation in the absence of IRAP for both types of stimulation (Fig. 1E).

The exocytosis of secretory granules depends on the SNARE VAMP8. We observed no or poor colocalization of IRAP with VAMP8 and preformed TNF-α stored in secretory granules in resting mast cells (Fig. 1F). Prior to release, granule-bound VAMP8 assembles with the t-SNAREs Stx4 and SNAP23 at the plasma membrane or in intracellular degranulation channels (Moon *et al*., 2014). Also during the degranulation process we failed to observe any colocalization of IRAP with VAMP8 (Fig. 1G) confirming the localization to distinct endosomal compartments.

However, we wondered if the increased release of granule content in IRAPko mast cells was reflected by an increased complex formation of VAMP8 with Stx4 upon activation. We therefore quantified colocalization of VAMP8-positive granules with Stx4 by imaging flow cytometry after stimulation with ionomycin/PMA. As expected, we observed more VAMP8 colocalization with Stx4 in IRAPko than in wild-type (wt) mast cells (Fig. 1H and Suppl. Fig.1). Next, we sought to assess degranulation *in vivo*. To this end, we challenged IRAP wt and ko mice on one ear for degranulation with arachidonic acid, while the other ear was left untreated. Arachidonic acid induces degranulation and cytokine production in mast cells through the prostaglandin EP3 receptor ^47^. In line with our *in vitro* results, we detected significantly more histamine in crude homogenates of stimulated IRAPko mouse ears than in wt ears (Fig. 1H). This was not due to different mast cell densities in tissues of wt compared to IRAPko mice (Suppl. Fig. 2).

To confirm the mast cell-specificity of this test, we reconstituted mast cell-deficient kit-W^sh/sh^ mice (Wsh) with bone marrow-derived mast cells (BMMC) from wt or IRAPko donor mice and challenged them along with non-reconstituted Wsh mice. As expected, no histamine was detected in the ear homogenates from the mast cell-deficient, non-reconstituted Wsh mice, while histamine secretion was increased in the challenged ears in Wsh mice reconstituted with IRAPko BMMC compared to wt BMMC (Fig. 1I). It is important to note that this assay does not discriminate the origin of the detected histamine from intracellular stores versus extracellular locations after degranulation. It is, however, likely that the histamine epitopes recognized in the antibody-based detection assay are more exposed after exocytosis, explaining the net increase of detectable histamine in challenged ears containing IRAPko mast cells, while smaller quantities of released molecules in wt ears might not be detectable with this protocol due to a strong background signal generated by histamine from intracellular stores.

In conclusion, we show that IRAP endosomes are dispensable for the VAMP8-dependent pathway of regulated secretion in mast cells, and moreover, in their absence, degranulation is increased *in vitro* and *in vivo*.

### Constitutive secretion of cytokines relies on IRAP endosomes in mast cells

Next to the regulated secretion of stored granule contents, mast cells produce and secrete *de novo* synthesized cytokines via the constitutive secretion pathway. Although both species of secreted vesicles originate from the Golgi and engage with Stx4 and SNAP23 for docking and fusion at the plasma membrane, they follow distinct post-Golgi trafficking routes. Thus, while regulated secretion depends on VAMP8, *de novo* synthesized cytokines in the constitutive pathway in murine mast cells stain with VAMP3 ^19^. We observed that IRAP endosomes colocalized well with VAMP3 in mast cells (Fig. 2A). VAMP3 is associated with Golgi trafficking to and from the recycling compartment and has been implicated in TNF-α secretion in macrophages ^21^. In mast cells, TNF-α is stored in limited amounts in secretory granules, and *de novo* produced and secreted via the constitute pathway in the late phase of activation. TNF-α is transported throughout the cell as a transmembrane pro-cytokine. Release of soluble TNF-α into the extracellular space requires the activity of the TNF-α-cleaving enzyme TACE. In the presence of the TACE inhibitor TAPI-I TNF-α accumulates at the surface of activated cells starting from 1h of activation, where it colocalized strongly with IRAP (Fig. 2B). We therefore hypothesized that IRAP might be involved in the constitutive secretion pathway of cytokines in mast cells.

**Fig. 2.**
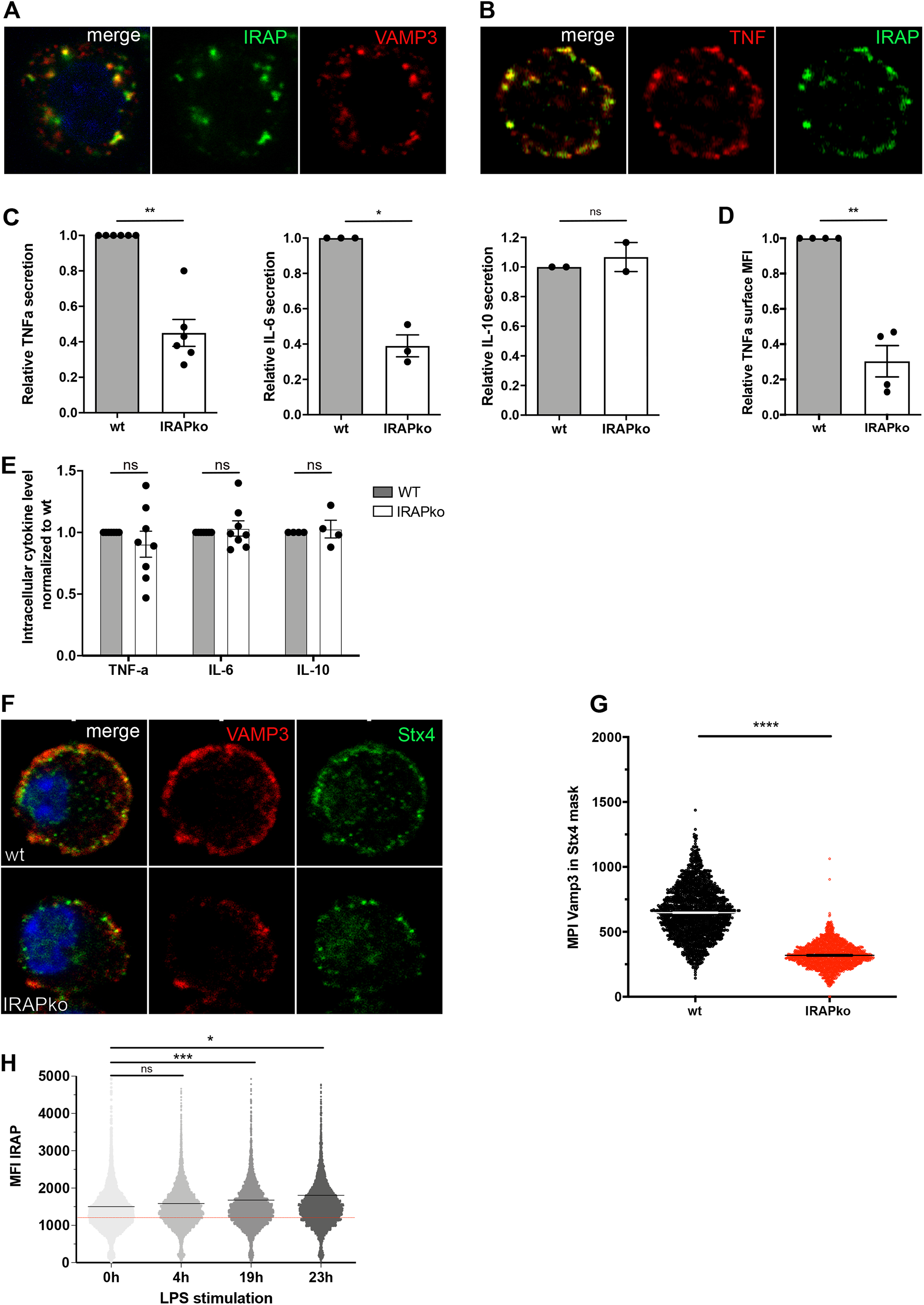
IRAP endosomes are required for pro-inflammatory cytokine secretion in mast cells. (A) Resting BMMCs were immunostained for IRAP and VAMP3. Images are representative of four independent experiments. (B) BMMCs were treated for 2h with ionomycin/PMA and TAPI-1, and immunostained for TNF-α and IRAP. Images are representative of three independent experiments. (C) Peritoneal mast cells were stimulated with ionomycin/PMA for 18h and secreted cytokines were quantified by ELISA in the culture supernatant. Graphs represent mean ± SEM of three or more experiments. (D) Peritoneal mast cells were stimulated for 4h with ionomycin/PMA and TAPI-I, and plasma membrane-bound TNF-α on live cells was detected via flow cytometry. Graphs represent mean ± SEM of four experiments. (E) Intracellular cytokine expression was determined by flow cytometry of fixed and permeabilized peritoneal mast cells after 4h of treatment with ionomycin/PMA and brefeldin A. Shown are mean values of IRAPko cells ± SEM of three experiments reported to wt. (F) BMMCs were stimulated with ionomycin/PMA for 1h prior to immunostaining for VAMP3 and Stx4 and confocal imaging. Images are representative of two independent experiments. (G) Imaging flow cytometry quantification of VAMP3 mean pixel intensity (MPI) in the Stx4 mask of ionomycin/PMA stimulated mast cells. Analysis was performed on >3000cells. (H) BMMC were exposed to LPS for indicated times and IRAP expression analyzed by flow cytometry in permeabilized cells. Red line indicates staining background of IRAPko cells. Graphs show mean fluorescence values ± SEM of >1000 cells. *p<0.05,** p<0.01, ***p<0.001, ns non significant

Indeed, TNF-α and IL-6 secretion was reduced about 50% in IRAPko compared to wt peritoneal mast cells after ionomycin/PMA stimulation as determined via ELISA (Fig. 2C) or TAPI-I treatment and TNF-α surface staining followed by flow cytometry analysis (Fig. 2D). Of note, secretion of the regulatory cytokine IL-10 was not affected by the absence of IRAP (Fig. 2C). In order to verify that the observed secretion defect was due to a trafficking defect in IRAPko cells rather than a diminished synthesis rate, we compared intracellular cytokine levels after 4h of activation under inhibition of Golgi/post-Golgi trafficking with brefeldin A. As we failed to detect any significant differences in the quantities of intracellularly produced cytokines between IRAP wt and ko mast cells under these conditions (Fig. 2E), we concluded that the absence of IRAP endosomes produces a trafficking defect of *de novo* synthesized IL-6 and TNF within or beyond the Golgi.

This defect also translates into reduced quantities of VAMP3-bearing vesicles colocalizing with Stx4 at the plasma membrane in IRAPko cells detectable by confocal imaging (Fig. 2F) and imaging flow cytometry (Fig. 2G).

Macrophages increase VAMP3 expression under LPS stimulation, possibly to cope with the need for more transport machinery upon increased cytokine synthesis ^21^. To test if the same was true for IRAP expression, we stimulated mast cells for different periods of time with LPS and measured IRAP expression by intracellular flow cytometry. Indeed, IRAP was induced over time (Fig. 2H), compatible with a role in pro-inflammatory cytokine trafficking.

### IRAP endosomes are required for TNF-α secretion *in vivo*

To quantify cytokine secretion by mast cells *in vivo*, we performed the mouse ear challenge experiment from above. While increased TNF-α and IL-6 levels were detected in wt ears within 45 min after the challenge, no cytokine release was observed in the ears of IRAPko animals (Fig. 3A and B). Confirming our *in vitro* results, IL-10 secretion was not affected by the lack of IRAP endosomes *in vivo* (Fig. 3C). Searching to monitor the mast-cell contribution to the observed effects, we repeated the test in Wsh mice that had been reconstituted with wt or IRAPko BMMC. Wsh mice reconstituted with IRAPko mast cells showed defects in TNF-α and IL-6 secretion indicating that the cytokines measured in this experimental setting could indeed be attributed to mast cells (Fig. D and E).

**Fig. 3.**
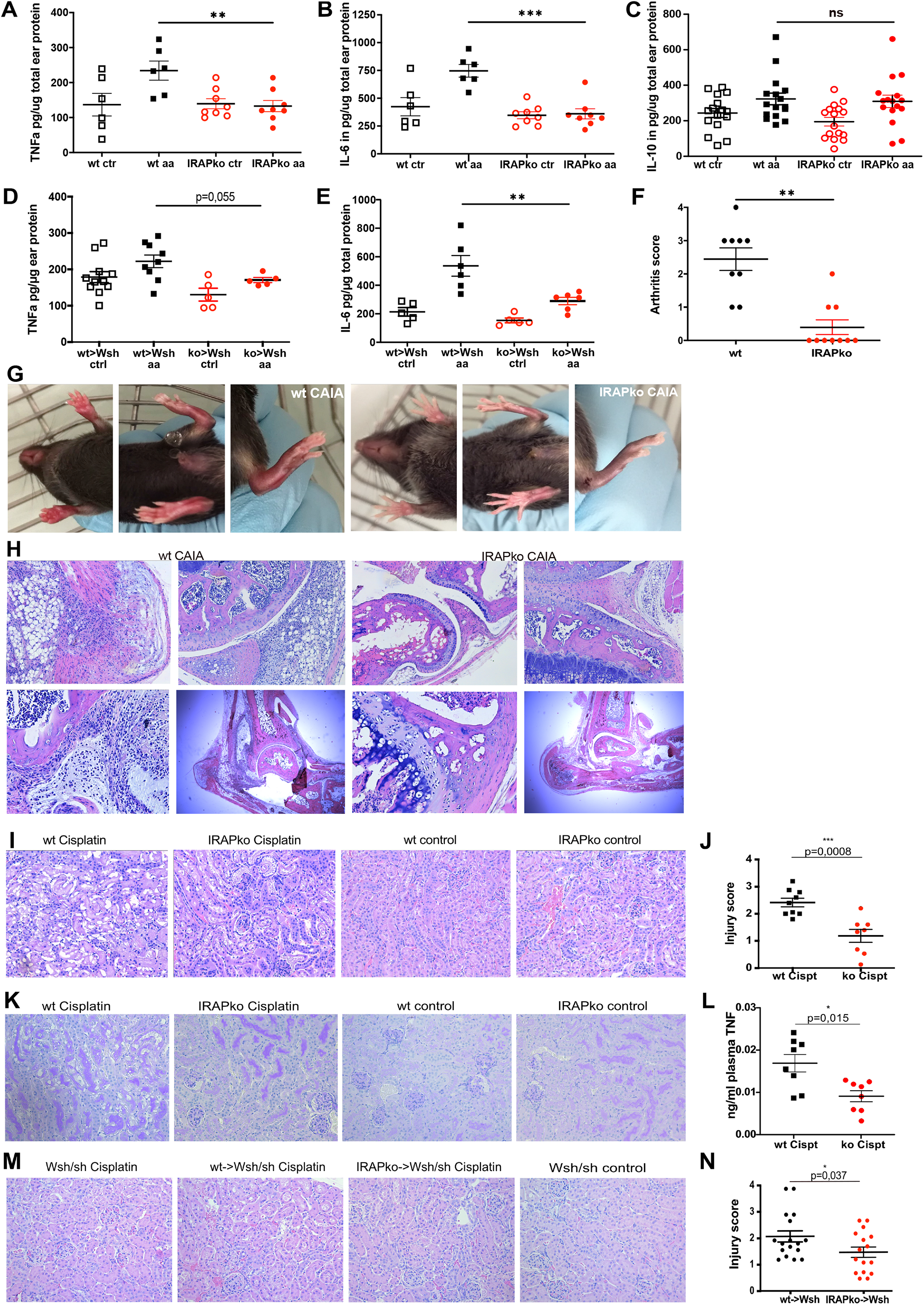
IRAP endosomes are required for inflammatory cytokine secretion *in vivo*. (A-E) IRAPwt and ko mice (A-C) or mast-cell deficient Wsh mice reconstituted with IRAPwt and ko BMMC (D,E) were challenged on one ear with 30mg/ml arachidonic acid while the control ear was left untreated. Cytokine concentrations were quantified in ear tissue homogenates and normalized to total protein concentration. (F) Arthritis score of IRAPwt and ko mice 8 days after CAIA induction. Graphs show pooled data from two independent experiments. (G) Photographs of front and hind limbs of representative IRAPwt and ko mice 8 days after CAIA induction. (H) Histological images of HE-stained paraffine sections of relevant hind limb joints of IRAPwt and ko mice at 8 days after CAIA induction. (I) Representative HE-stained paraffin sections of cisplatin-treated and control IRAPwt and ko mouse kidneys. (J) Kidney injury score of cisplatin-treated IRAPwt and ko mice. (K) Periodic acid Schiff-stained paraffin kidney sections of cisplatin-treated and control IRAPwt and ko mice. (L) Plasma TNF-α concentration of cisplatin-treated IRAPwt and ko mice 24h after cisplatin injection from one out of three experiments. (M) HE-stained paraffin kidney sections of cisplatin-treated non-reconstituted and IRAPwt or ko BMMC-reconstituted kit-Wsh/sh mice from one out of three experiments. (N) Injury scores of paraffin kidney sections of cisplatin-treated IRAPwt or ko BMMC-reconstituted kit-Wsh/sh mice from three independent experiments. *p<0.05,** p<0.01, ***p<0.001

The role of TNF-α in the pathogenesis of collagen-induced arthritis (CAIA) is well documented ^48,49^. To examine the relevance of IRAP endosomes for TNF-α secretion in this disease model, we challenged wt and IRAPko mice with an arthritogenic collagen-directed antibody cocktail. Eight days after arthritis induction, wt mice presented signs of joint inflammation marked by intense redness, swelling of paws and joints and difficulties to walk, while the majority of IRAPko mice showed no or only mild symptoms (Fig. 3F, G). Histological analyses of the knee (3H, upper panels) and ankles (3H, lower panels, right) revealed joint swelling accompanied by strong infiltration of inflammatory cells into the synovial space and bone erosion in wt animals. IRAPko mice showed less or no infiltrations, less swelling and no bone damage. We conclude that IRAP is required for the strong inflammatory disease phenotype observed in wt mice.

As the role of mast cells in this model has been somewhat questioned by the fact that Kit-W^v/v^ but not Wsh mice were protected from CAIA ^13,50^, presumably due to differences in their megakaryocyte populations ^51^, we turned to a cisplatin-induced kidney inflammation model previously reported to depend on mast cell-derived TNF-α as an alternative approach ^12^. Cisplatin is an efficient and widely employed cytostatic agent for cancer therapy the tolerance for which, however, is limited by the frequent adverse effect of acute kidney injury. We hypothesized that the TNF-α-dependent kidney injury after cisplatin administration that is characterized by tubular apoptosis, necrosis and inflammation, would be attenuated in IRAPko mice. To verify this, we histologically analyzed the kidneys of IRAP wt and ko mice 96h after peritoneal cisplatin injection. HE (Fig. 3I) and PAS (Fig. 3K) staining of paraffin-embedded kidney samples revealed visibly reduced tubular damage in IRAPko animals and translated into significantly lower injury scores that were determined independently in a blinded evaluation by three different experimenters (Fig. 3J). These observations were consistent with significantly reduced TNF-α plasma levels in cisplatin-treated IRAPko mice (Fig. 3L). Cisplatin-induced inflammation and nephrotoxicity are mediated via TLR4 ^52^. In order to exclude the possibility that different TLR4 expression levels of wt versus IRAPko cells were at the origin of the observed effects, we confirmed comparable TLR4 surface expression of wt and IRAPko mast cells (Suppl. Fig. 3)

To evaluate the contribution of mast cells to these effects, we administered cisplatin to wt or IRAPko BMMC-reconstituted Wsh mice. Scoring of the histological injury level indicated that the mean kidney damage of mice reconstituted with IRAPko mast cells was reduced as compared to mice reconstituted with wt mast cells (Fig. 3M, N), although the difference was less pronounced than between wt versus ubiquitously IRAPko mice. The involvement of other TNF-α-producing cell types in the tested cisplatin model might explain the limited differences in the experiments with mast cell-reconstituted mice. We therefore addressed TNF-α secretion in peritoneal macrophages using confocal imaging and the TAPI-based flow cytometry assay described above. IRAP colocalized strongly with TNF-α in ionomycin-activated macrophages (Suppl. Fig. 4A). Furthermore, IRAPko macrophages showed diminished TNF surface staining after 4 hours of activation by ionomycin/PMA or LPS (Suppl. Fig. 4B). We conclude that also macrophages depend on IRAP endosomes for the efficient secretion of TNF-α via the constitutive pathway.

Taken together, IRAPko mast cells, and likely other immune cell types, secrete less TNF-α *in vivo* leading to milder phenotypes in TNF-α -dependent murine disease models.

### IRAP is required for Golgi export of TNF-α transport vesicles

We next sought to unravel at which step the exocytic cytokine trafficking was impaired in the absence of IRAP. To this end we analyzed Stx6 colocalization with VAMP3 in activated mast cells. Stx6 decorates IRAP vesicles in different cell types and is present on TNF-α carriers after budding from the Golgi in macrophages ^21,22^. While Stx6 colocalized well with VAMP3 at the plasma membrane in activated mast cells, significantly less Stx6 was detected in the VAMP3-stained areas in IRAPko cells due to overall reduced peripheral Stx6 staining (Fig. 4A and B). Total VAMP3 levels are also reduced in IRAPko mast cells (Suppl. Fig. 5). These observations prompted us to inquire if TNF-α carriers required IRAP for budding from the Golgi. We therefore adapted a previously published Golgi export assay ^53^. LPS-pre-activated cells were incubated at 20°C for 3h to enrich cytokines in the Golgi. Subsequent temperature shift to 37°C re-activates budding of exocytic transport vesicles from the Golgi allowing for analysis of Golgi export kinetics of cytokines in the constitutive pathway.

**Fig. 4.**
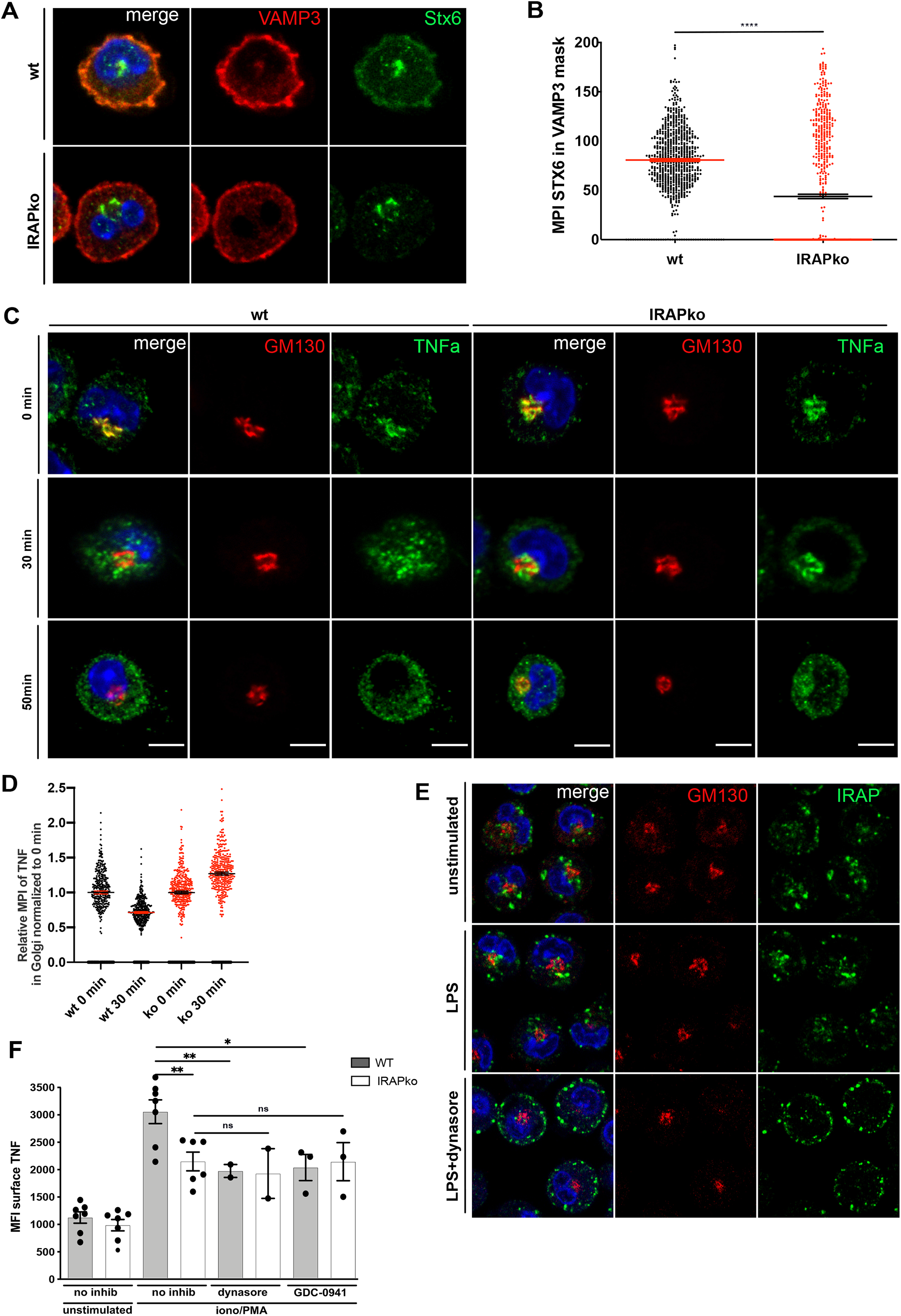
IRAP/Stx6 endosomes are required for formation of VAMP3+ endosomes for TNF-α secretion. (A) Representative confocal images of VAMP3 and Stx6 of LPS-stimulated mast cells. (B) Imaging flow cytometry quantification of Stx6 staining in VAMP3 mask of LPS-stimulated mast cells. Graphs show mean fluorescence intensities ± SEM of >700 cells. (C) BMMC were stimulated with LPS for 18h at 37°C, 5% CO_2_, placed at 20°C for 3h, and either fixed (0min) or re-incubated at 37°C for 30min or 50min before staining for GM130 and TNF-α. Representative confocal images of IRAPwt and ko cells from one out of four independent experiments. (D) Imaging flow cytometry quantification of TNF-α Golgi export kinetics. Graphs show mean pixel intensities of TNF-α in the GM130 mask reported to the 0min sample from >3000 cells. (E) Mast cells were activated with 100ng/ml LPS and 80uM dynasore was added after 60min. Cells were fixed and stained for IRAP and GM130 after 4h. (F) Peritoneal mast cells were treated with TAPI-I and ionomycin/PMA. Dynasore (80uM) and GDC-0941 (1.5uM) were added after 60min and TNF-α was detected via surface FACS staining after 4h of activation. Results are presented as means ± SEM from three (two for dynasore) independent experiments. *p<0.05,** p<0.01, ***p<0.001

After 3h at 20°C, TNF-α colocalized strongly with the Golgi marker GM130 in both wt and IRAP ko cells (Fig. 4F, 0 min time point), indicating successful inhibition of Golgi export under these conditions. Re-activation of exocytic trafficking resulted in progressive export of TNF-α from the Golgi in wt cells, while in IRAPko cells, a net accumulation was observed over the first 30min, indicating that the translation rate exceeded the export rate in these cells (Fig. 4E). At 50min, IRAP expressing cells had largely emptied the Golgi of TNF-α, while in IRAPko cells, colocalization of TNF-α and GM130 persisted (Fig. 4F).

Considering that the cytosolic retention pool of IRAP vesicles, upon activation, has been proposed to translocate to the plasma membrane without passing through the Golgi ^38,54^, intersection with TNF-α carriers is difficult to envisage. However, under prolonged activating signaling, IRAP is reinternalized and retrieved to the Golgi from sorting endosomes via retromer action ^36^. We therefore wondered if endocytosis inhibition changed the subcellular localization of IRAP and ultimately TNF-α secretion. Indeed, in the presence of the dynamin inhibitor dynasore IRAP showed a strong plasma membrane staining (Fig. 4E) after three hours of LPS activation indicating efficient inhibition of IRAP re-internalization. Consistently, both dynasore as well as the PI3K I inhibitor GDC-0941 were able to reduce TNF-α secretion specifically in wt cells (Fig. 4F). These results suggest that IRAP internalization is a prerequisite for normal TNF-α trafficking (Suppl. Fig. 6).

### IRAP inhibition by HFI-419 destabilizes IRAP endosomes

Considering the aminopeptidase function of IRAP, we wondered if its catalytic activity was required for efficient cytokine secretion. To test this, we treated mast cells with the IRAP inhibitors HFI-419, 4u ^55^ and 22b (a gift from E. Stratikos) for 24h prior to ionomycin/PMA activation. Although all three inhibitors showed a tendency to inhibit IRAP-dependent cytokine secretion *in vitro*, only HFI-419 mediated significant inhibition (Fig. 5A) and was therefore selected for further *in vivo* studies.

**Fig. 5.**
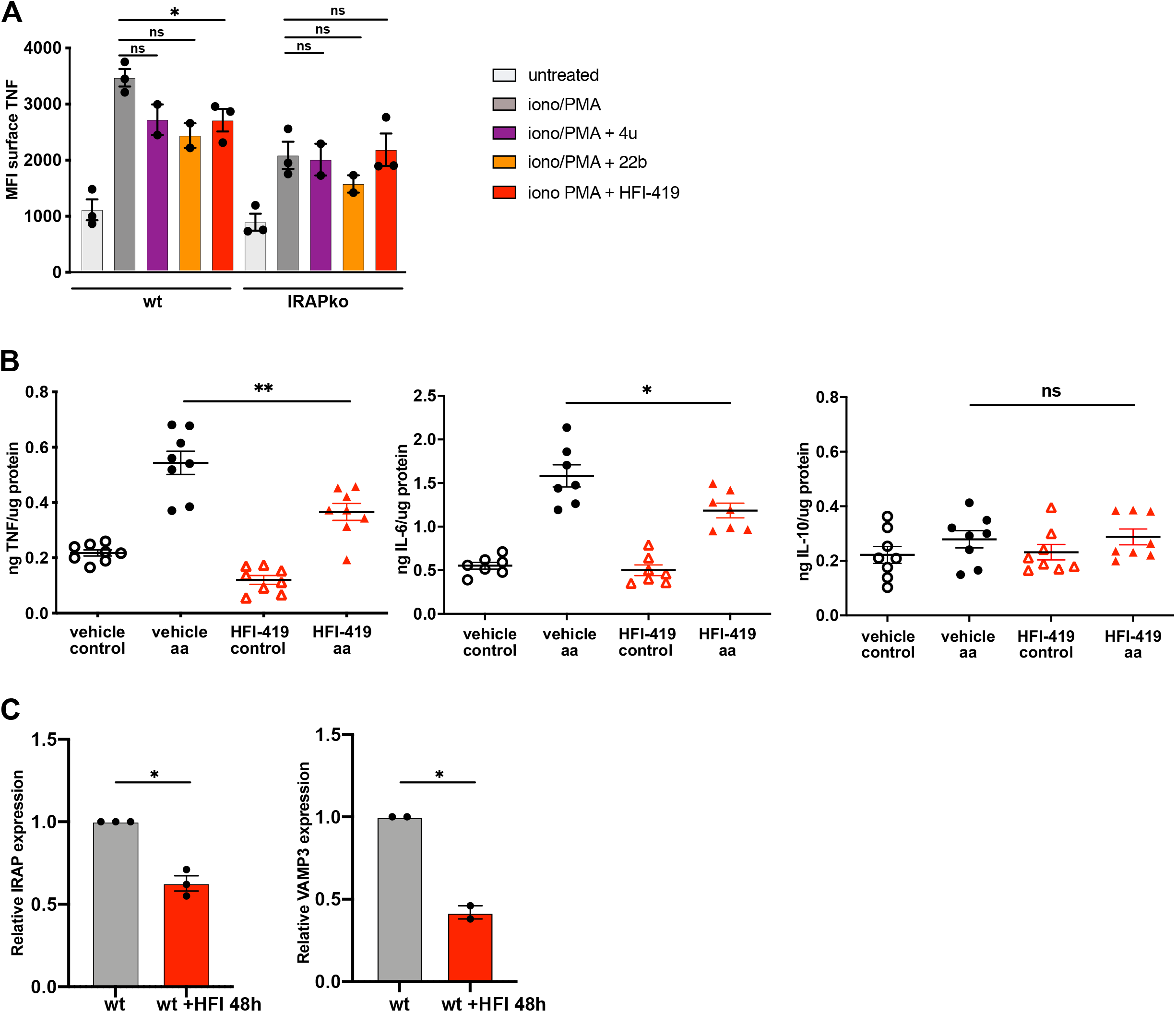
IRAP inhibitor HFI-419 blocks cytokine secretion via destabilization of IRAP and VAMP3+ endosomes. (A) Peritoneal mast cells were cultured in the presence or absence of different IRAP inhibitors (1µM each) for 24h and exposed to TAPI-I and ionomycin/PMA for 4h or left untreated. Cell surface-bound TNF-α was detected via flow cytometry in unpermeabilized live cells. Graphs show means ± SEM of three (two for 4u and 22b) independent experiments. (B) IRAP wt mice were injected i.v. with 6µg HFI-419 or vehicle 24h and 15min prior to ear challenge. Cytokine concentrations were quantified in ear tissue homogenates and normalized to total protein concentration. Graph shows one representative out two experiments. (C) BMMC were exposed to the IRAP inhibitor HFI-419 for 24h, and IRAP and VAMP3 expression determined via intracellular staining in flow cytometry. Graphs show means ± SEM of three (IRAP) or two (VAMP3) experiments.

Vehicle or inhibitor at a dose of 1µmol/kg was administered intravenously at 24h and 15min prior to the ear challenge. HFI-419-treated animals secreted significantly less TNF-α and IL-6 than vehicle-treated animals, while IL-10 secretion was unaffected, indicating that the availability of the catalytic domain of IRAP was directly or indirectly required for trafficking of these pro-inflammatory cytokines. This was somewhat surprising with respect to previous studies in different cell types in which reconstitution by a protease-dead IRAP variant fully restored vesicular distribution and endosomal trafficking in IRAPko cells, suggesting that the enzymatic activity was dispensable for IRAP-mediated trafficking functions ^43,45^. We therefore hypothesized that in addition to blocking its catalytic activity, HFI-419 may also affect the stability of IRAP. In agreement with this hypothesis, using intracellular flow cytometry staining, we detected decreased expression levels of IRAP and VAMP3 starting from 24h of HFI-419 treatment (Fig. 5C and Suppl. Fig. 7), suggesting that ligation of the inhibitor HFI-419 induced IRAP degradation, ultimately hampering TNF-α secretion.

## Discussion

In the present study we demonstrate that IRAP controls late-phase pro-inflammatory cytokine secretion in mast cells *in vitro* and *in vivo*. We find that in Ca^2+^-activated IRAPko mast cells, secretion of TNF-α and IL-6 was reduced compared to wt cells. This was due to a trafficking defect rather than reduced cytokine synthesis because intracellular cytokine levels were comparable between wt and IRAPko cells in response to intracellular Ca^2+^ triggers. The observed inhibition of cytokine secretion was in the order of 50%, and this reduction was physiologically relevant, as IRAPko mice showed milder disease phenotypes in two experimental models of TNF-α-dependent pathologies, namely CAIA and cisplatin-induced acute kidney injury. Moreover, human GWAS data analysis revealed that a variant of the Lnpep gene (rs27293) is associated with significantly increased risk of developing juvenile idiopathic arthritis, a disease with established TNF-α involvement. Considering that expression of the minor (risk) allele results in higher IRAP mRNA levels than the major (protective) allele, these data corroborate our findings on IRAP in TNF-α secretion ^56^. Previous reports have implicated the recycling endosome-related SNARE VAMP3 in the constitutive secretion pathway in mast cells ^17,19^ and macrophages ^21^. We extend these findings by showing that, in the absence of IRAP, the amount of VAMP3 colocalizing with Stx4, the SNARE involved in vesicle fusion with the plasma membrane, was reduced. This was most likely due to the observed reduction in the formation or stabilization of Stx6+ post-Golgi carriers in IRAPko cells.

Consistently, a Golgi export assay confirmed that, in IRAPko mast cells, *de novo* synthesized TNF-α persisted in the Golgi for longer periods of time. These results collectively hint to a role for IRAP in formation of Stx6+ carriers in charge of TNF-α and IL-6 transport at the TGN. The prevailing localization of IRAP to a sequestered pool of cytoplasmic vesicles in the steady-state is difficult to reconcile with a role as sorting effector at the TGN. However, IRAP vesicles are mobilized in response to specific activation signals which induce cleavage of the cytosolic retention protein TUG and transport of IRAP to the cell surface ^34^. Importantly, it has been suggested that under prolonged stimulation, IRAP recycles through endosomes and Golgi back to the plasma membrane without transit via the retention pool ^38,54^. In the present study, we identify this exocytic/recycling pathway of IRAP as overlapping with constitutive cytokine secretion. Moreover, the defective TNF-α secretion in the presence of endocytosis inhibitors that was specifically observed in IRAPwt cells suggests that IRAP endocytosis is required for efficient post-Golgi trafficking of cytokines. Taken together, we suggest that activation of mast cells results in IRAP mobilization to the plasma membrane, re-internalization and retrieval to the TGN where it functions as a sorting receptor for cytokines and possibly other molecules secreted along the constitutive pathway.

Post-Golgi transport vesicles containing TNF-α and IL-6 are formed through fission of tubular compartments from the TGN. Budding of these tubular carriers occurs from different TGN subdomains and depends on different coiled-coil golgins ^57^. For instance, the transporters involved in TNF-α exit from the Golgi are positive for golgin-245/p230 ^58^, while the sorting and export of Glut4 and IRAP to the sequestered GSV pool in adipocytes is golgin-160 dependent. Importantly, upon depletion of golgin-160, Glut4 is routed to the PM ^59^. We therefore speculate that under persistent activation, the interaction between IRAP and golgin-160 is abrogated, possibly through a post-translational modification of IRAP, changing the post-Golgi trafficking of IRAP and sorting it into a distinct, most likely golgin-245-dependent pathway to the PM.

Interestingly, IL-10 secretion was not affected by the loss of IRAP. Studies in macrophages, where the secretion pathways of TNF-α, IL-6 and IL-10 have been studied in detail, revealed that while these three cytokines may use a common route from the TGN to the recycling endosome, IL-10 alternatively uses a distinct post-Golgi pathway that overlaps with trafficking of the lipoprotein ApoE ^60^. Based on our results we suggest that, at least in mast cells, the portion of IL-10 trafficking along the same IRAP-dependent pathway as IL-6 and TNF-α is minor.

With respect to regulated exocytosis, we observed increased secretory granule release in mast cells in the absence of IRAP. Consistently, more VAMP8 staining was observed on Stx4-positive membrane domains, indicating increased fusion events between secretory granules and the plasma membrane in IRAPko as compared to wt cells.

This dichotomy of impaired constitutive trafficking and augmented secretory lysosome/granule trafficking is reminiscent of a report on sortilin ko cytotoxic T and NK cells. In these cells, sortilin has been suggested to regulate both VAMP7 targeting to lysosomes and constitutive secretion of IFNlll (but not TNF-α) ^53^.

Although a direct role for IRAP in the lysosomal targeting or degradation of Vamp8 seems unlikely considering the absence of colocalization between those two proteins, we cannot exclude IRAP-dependent trafficking of proteins that negatively regulate VAMP8 degradation. Alternatively, considering that the same SNARE docking and fusion machinery is used for exocytosis of VAMP3+ vesicles and VAMP8+ granules, diminished abundance of VAMP3+ carriers at the PM might leave more Stx4-SNAP23 molecules available for SNARE complex formation with VAMP8, ultimately augmenting the VAMP8-dependent degranulation rate. Moreover, the activity of several VAMP family members including VAMP8 can be regulated via phosphorylation through PKC which terminates the degranulation response ^61^. This regulation likely prevents dangerous consequences of excessive degranulation from mast cells, notably anaphylactic shock. In contrast, this level of regulation is lacking for VAMP3 due to the absence of a phosphorylation motif ^61^, suggesting that other regulatory mechanisms may exist. The implication of signal-responsive IRAP endosomes in VAMP3-dependent exocytosis might constitute such a mechanism, i.e. linking extracellular cues to cytokine trafficking.

IRAP protein expression was induced by LPS, in agreement with a previous report that showed IRAP mRNA induction by LPS and IFN-⍰ but not TGF-β in macrophages ^62^. These findings suggest that IRAP endosomes are part of a transcriptionally regulated trafficking machinery that is induced by pro-inflammatory environmental cues. Particularly in the light of a recently emerging polarization concept for mast cell functions in inflammation and cancer, in analogy to macrophage M1 vs M2 polarization, the transcriptional regulation of IRAP endosomes deserves further exploration.

We also showed that macrophages depend on IRAP expression for TNF secretion. Considering the broad expression profile of IRAP amongst immune cells, IRAP might regulate cytokine secretion in other cell types, especially those that need to maintain a temporal or spatial segregation between regulated secretion of stored granules and constitutive secretion, such as platelets, cytotoxic T cells, NK cells and basophils.

Finally, IRAP expression was reduced using the chemical inhibitor HFI-419. Considering that HFI-419 binds to the substrate binding pocket in the intraluminal region of IRAP ^63^, the diminution in protein levels strongly suggest conformational effects in trans acting on the cytosolic tail of IRAP ^64^, which contains specific motifs for the regulated trafficking and interaction of IRAP with several proteins involved in vesicular trafficking such as formins ^44,65^, tankyrase ^66^ and p115 ^67^. We have previously shown that the loss of IRAP anchoring to the actin cytoskeleton promoted destabilization and degradation of IRAP endosomes through rapid retrograde dynein-mediated transport and fusion with lysosomes ^42,44^.

In summary, our results identify IRAP as a transcriptionally regulated hub of late phase cytokine secretion in mast cells and a potential target for anti-inflammatory drug development.

## Methods

### Reagents and antibodies

The following antibodies were used in this study. Mouse monoclonal IRAP antibody clone 3E, rabbit monoclonal anti-IRAP XP clone D7C5, rabbit anti-EEA1 (all Cell Signaling); rat anti-mouse lysosome-associated membrane protein (LAMP)1 clone 1D4B, mouse monoclonal anti-STX6, mouse monoclonal anti-GM130 (BD Pharmingen); rabbit polyclonal anti-STX6 (ProteinTech Group, Chicago, IL, United States); mouse monoclonal anti-Stx4 clone QQ-17 (Santa Cruz); rabbit polyclonal anti-TNF (abcam 34674 for confocal imaging and imaging flow cytometry), PE-PerCP5.5 anti-mouse TNF clone MP6-XT22 (eBiosciences for FACS); rat anti-IL-6 and rat anti-IL-10 (eBiosciences); rabbit polyclonal anti-VAMP3 (abcam 2102); rabbit polyclonal anti-VAMP8 (novus); goat anti-serotonin (abcam 66047), rabbit polyclonal anti-Rab14 (Sigma Aldrich). All secondary reagents were Alexa-coupled highly cross-adsorbed antibodies from Molecular Probes (Life technologies). Alexa647-transferrin was from Life technologies. IL-3 and SCF (premium grade) were purchased from Milteny Biotec.

Murine cytokine detection Duoset enzyme-linked immunosorbent assay (ELISA) kits were from R&D Systems (mIL-6, mTNF) or from Biolegends (mIL-10). Easysep™ anti-mouse CD117 positive selection kit was from Stemcell. The TNF-alpha–converting enzyme (TACE) inhibitor TAPI-1, ionomycin, PMA, HFI-419, dynasore, GDC-0941 were all from Calbiochem. p-Nitrophenyl-N-acetyl-β-D-glucosaminide (pNAG) was from Sigma. The IRAP inhibitors 4u and 11b were a gift from E.Stratikos (Demokritos Research Center Athens).

### Mice

Previously described IRAP^−/–^ mice on an Sv129 background obtained from S. Keller were back-crossed up to 10 times to C57BL/6 mice obtained from Janvier (St. Quentin-Fallavier, France). Control wt mice were C57BL/6 mice bred in our facility or C57BL/6 mice purchased from Charles River. Kit-W^sh/sh^ were purchased from Jackson Laboratories (strain #30764).

### Mast cell isolation and culture

Murine bone marrow-derived mast cells (BMMCs) were produced *in vitro* by culturing cells extruded from large bones for 4 to 6 weeks in complete medium [Iscove’s modified Dulbecco’s medium (IMDM) complemented with 10% fetal calf serum (FCS), 25 mM HEPES (pH 7.4), 2 mM glutamine, 100 U/ml penicillin, 100 g/ml streptomycin, 50 µM β-mercaptoethanol, 1% non-essential amino acids and 1mM sodium pyruvate] supplemented with 10ng/ml IL-3.

Every 5-7 days, medium was replaced. All cell cultures were grown at 37°C in a humidified atmosphere with 5% CO2. BMMC differentiation as verified by staining with CD117 and FcεR antibodies after 4 weeks was more than 98%. Mouse peritoneal-derived mast cells (PCMCs) were obtained by peritoneal lavage with 5 mL ice-cold PBS/0.1% BSA and cultured in complete medium (see above) supplemented with 10ng/ml IL-3 and 10ng/ml SCF.

Non-adherent cells including mast cells were separated from adherent macrophages after 3h of culture. Cultured cells were enriched for mast cells (>90%) after 7 days of culture. For use after shorter culture times, mast cells were purified via anti-CD117 beads (StemCell).

### Mast cell reconstitution of kit-W^sh/sh^ mice

BMMC from wt and IRAPko mice were cultured for 4 weeks in the presence of murine IL-3 and murine SCF as described above. 5×10^6^ BMMC were injected i.v. in kit-W^sh/sh^ mice and allowed for 8 to 12 weeks for reconstitution before functional experiments. Reconstituted mice yielding less than 50nM histamine per µg total protein in untreated ear tissue homogenates were considered as unsuccessfully reconstituted and excluded from the analysis.

### Flow cytometric assays

For TNF-α surface staining, PCMCs were stimulated with 1µM ionomycin/10nM PMA or 100ng/ml LPS at 37°C for 4h in the presence of TAPI-1, washed with ice-cold PBS, and incubated at 4°C with Fcblock (Miltenyi) followed by fluorochrome-conjugated CD117, FcεRI and TNF-α antibodies diluted in PBS-1% BSA. Intracellular staining of cytokines, IRAP and VAMP3 was performed using the BD intracellular staining kit and suitable species-specific fluorescent secondary antibodies (Life technologies).

For degranulation experiments, PCMCs were stimulated with 1µM ionomycin/10nM PMA or 48/80 for 30 min at 37°C, placed on ice and surface-stained with AlexaFluor488 anti-LAMP1. BD Canto™ and Gallios flow cytometers were used for cell analysis.

### Mouse ear challenge

Twenty microliters arachidonic acid (AA) (30mg/ml in acetone) were applied to the inner and outer surface of one mouse ear, whereas the other ear was left untreated.

One hour after AA application, mice were sacrificed and ears were collected. The ear biopsies were dissociated using the pre-set “Protein” protocol of gentleMACS™ Octo Dissociator (Milteniy Biotec) in 800µl ice-cold homogenization buffer [(PBS containing 0,4M NaCl, 0,05% Tween-20, 10mM EDTA and protease inhibitor cocktail complete (Roche)]. The homogenates were cleared by 10 min centrifugation at 5000 x g, and the total protein concentration determined in a BCA assay. Histamine or cytokines in the supernatant were quantified as described below.

### Cytokine and histamine measurements

PCMCs were stimulated with 1uM ionomycin/10nM PMA or 100ng/ml LPS at 37°C for 6h for cytokine secretion or with ionomycin/PMA or 10ug/ml 48/80 for 30min for histamine measurement.

Supernatants were collected and histamine was quantified using the Histamine Dynamic HTRF kit (Cisbio). TNF-α, IL-6, or IL-10 were quantified using specific cytokine ELISA kits. Kits were used according to the manufacturer’s instructions.

### Beta-hexosaminidase release

PCMCs were stimulated with 1µM ionomycin/10nM PMA or 10ug/ml 48/80 for 30min in Tyrode’s buffer. Following stimulation, cell suspensions were centrifuged, placed on ice and supernatants were collected. The cell pellets of unstimulated cells were lysed with 0.5% Triton X-100 to determine the maximal enzymatic activity of β-hexosaminidase. Twenty-five microliters of supernatant or the lysate volume corresponding to 5×10^3^ cells were incubated with 50µl of a 1.3mg/ml p-Nitrophenyl-N-acetyl-β-D-glucosaminide (pNAG) solution in 50mM citrate buffer pH 4.5 at 37°C for 90min. The reaction was stopped with 150ul of 0.2 M glycine buffer pH 10.7 and absorbance was read at 405 nm. The percentage of degranulation was expressed as the ratio of absorbance of a given supernatant to the absorbance measured in the lysates of unstimulated cells.

### Cisplatin-induced kidney injury model

Mice were injected intraperitoneally with 10mg/kg cisplatin. Blood samples for measurement of plasma TNF-α levels were taken at 24h after cisplatin injection. Mice were sacrificed at 96h, and kidneys were processed for histological analysis as described in the histology section below. Tubular injury was independently scored in a blinded manner by three investigators.

### Collagen-antibody induced arthritis model

Mice were injected intravenously with 4mg/mouse antibody cocktail to collagen II (Chondrex, Inc.) on day 0, followed by an intraperitoneal LPS injection (25ug/mouse) on day 3. Severity of arthritis was evaluated on day 8 according to a qualitative scoring system as followed: 0 - normal, 1 - mild but definite redness of the ankle or wrist, or apparent redness and swelling limited to individual digits, 2 – moderate redness and swelling of ankle or wrist, 3 - severe redness and swelling of the entire paw including digits, 4 – maximally inflamed limb involving multiple joints. Mice were sacrificed and hind legs were collected, and processed for histological analysis as described below.

### Histology

Mouse ears, kidneys or hind legs were collected and fixed in 10% formalin for 24h. Hind legs were decalcified in 1M EDTA solution for 1 week. After paraffin embedding, 4µm sagittal (legs), transversal (ears) or coronal sections (kidneys) were cut and stained with hematoxylin/eosin or periodic acid Schiff stain as indicated. Tissue sections were imaged using a Leica DM2000 microscope equipped with a MC160HD camera using 5x and 20x objectives.

### Confocal microscopy

BMMCs were seeded on IBIDI poly-lysin-coated microscopy chambers in complete medium containing IL-3 at 37°C in a humidified atmosphere with 5% CO2 for 16h, stimulated as indicated, washed in PBS and fixed in PBS-4%PFA for 15min at room temperature. Permeabilization, blocking, washes and antibody incubation were performed in PBS-0.1% saponin/ 0.2% BSA at 18°C. Image acquisition was performed on a Zeiss LSM700 with an 63x oil-immersion objective. Images were analyzed and assembled using FIJI with the FigureJ plug-in.

### Imaging flow cytometry

One million BMMCs were stimulated as indicated, fixed with 4% PFA for 10min, permeabilized with permeabilization buffer (Invitrogen) and stained for indicated markers for 30min at RT, followed by a washing step in permeabilization buffer and secondary staining with fluorescently labeled antibodies for 30min at RT. Cells were washed, resuspended in PBS-2% FCS. Image acquisition was performed at 60X magnification using an ImageStream XMkII multispectral imaging flow cytometer (Amnis Corp., Seattle, USA), and acquired images were analyzed with the IDEAS software (version 6.2; Amnis Corp.). For SNARE protein analysis, a Stx4+ mask was defined and the mean pixel intensity of VAMP8 or VAMP3 was measured inside the mask. In the Golgi export assay, the Golgi mask was defined by GM130 staining and TNF mean pixel intensity quantified within the Golgi mask. Statistical analysis

Values are expressed as mean ± SEM, unless otherwise specified. Statistical significance between two groups was analyzed using the unpaired t-test with Welch’s correction, or one-sample t-test where replicates were expressed as percentage of a control group. P values are indicated as: * p < 0.05; ** p < 0.01; *** p < 0.001; **** p < 0.0001, ns = non-significant. GraphPad Prism version 9.0 was used to perform the statistical analysis.

### Study approval

Animal experimentation was conducted in agreement with the guidelines of local authorities, approved by the review board *Comité d’Éthique pour l’Expérimentation Animale* of Paris Descartes University.

## Author contributions

MW designed, performed, analyzed and interpreted research and wrote the manuscript. RR, CC and EE designed and performed experiments, MD performed and analyzed imaging flow cytometry experiments, RD performed IRAP inhibitor experiments. PvE, TTM and OH acquired funding, designed and interpreted research and co-wrote the manuscript.

## Supplementary Figures

**Suppl. Fig.1 VAMP8 – Stx4 co-staining in resting and activated mast cells**

(A) Representative confocal images of VAMP8 and Stx4 of resting wt and IRAP-deficient BMMC.

(B) Representative confocal images of VAMP8 and Stx4 of 10 min ionomycin/PMA-activated wt and IRAP-deficient BMMC.

**Suppl. Fig.2 Mast cell frequencies in IRAPko and wt mice**

(A) Toluidine blue staining of paraffin tissue samples and mast cell numbers

(B) Toluidine blue staining of paraffin paw tissue samples and mast cell numbers

(C) Flow cytometry staining of peritoneal cells and quantification of mast cell frequency

**Suppl. Fig.3 TLR4 surface levels of wt and IRAP deficient mast cells**

Flow cytometry staining with PE anti-TLR4 (clone MTS510, Sony Biotechnology Inc.) in resting c-kit+-gated BMMC.

**Suppl. Fig.4 TNF-α secretion in macrophages**

(A) Confocal images of TNF-α and IRAP-co-stained peritoneal macrophages exposed to ionomycin/PMA and TAPI-1 for 1h.

(B) Peritoneal macrophages were stimulated with LPS or ionomycin/PMA in the presence of TAPI-1 for 3h or left untreated and surface-stained with anti-TNF-α for analysis via flow cytometry.

**Suppl. Fig.5 VAMP3 in wt and IRAPko mast cells**

Quantification of VAMP3 staining of >3000 wt and IRAPko BMMCs by imaging flow cytometry.

**Suppl. Fig.6 Proposed trafficking scheme of IRAP and TNF in mast cells**

In resting cells, IRAP is sequestered to a cytoplasmic pool of small vesicles close to the Golgi. Upon activation via cell surface receptors resulting in intracellular Ca^2+^ level elevation, IRAP is rapidly translocated to the cell surface and then re-internalized into early endosomes from where it reaches the sorting endosome. From here it returns to the cell surface via recycling endosomes, or is transported to the Golgi through retromer binding, from where it employs the constitutive secretion pathway together with TNF-α. In IRAPko cells, or upon IRAP inhibitor treatment, TNF-α is blocked in the Golgi, while degranulation of secretory granules is facilitated. EE early endosome, SE sorting endosome, RE recycling endosome, GSV/IRV Glut4 storage vesicle/insulin-responsive vesicle, MVB multivesicular body.

**Suppl. Fig.7 IRAP inhibitor effect on IRAP expression**

IRAPwt and ko BMMC were cultured in the presence of 1µM HFI-419 for indicated times before fixing and intracellular staining for flow cytometry.

## Acknowledgments

The authors are grateful for the technical support of the Imaging Platform and the Histology Platform of the Necker Research Campus. This work was supported by State funding from the Agence Nationale de la Recherche (ANR) under “Investissements d’avenir” program (ANR-10-IAHU-01), “Blanc” programs ENDOREG (ANR-14-CE11-0014) and TIEBET (ANR-16-CE14-0024-01). Olivier Hermine’s laboratory is supported by Ligue Contre le Cancer (EL2018-HERMINE).

