## Supplementary figures and images for "Mast cell-mediated inflammation relies on insulin-regulated aminopeptidase controlling cytokine export from the Golgi"

### Supplemental Figure 1

**Suppl. Fig.1**

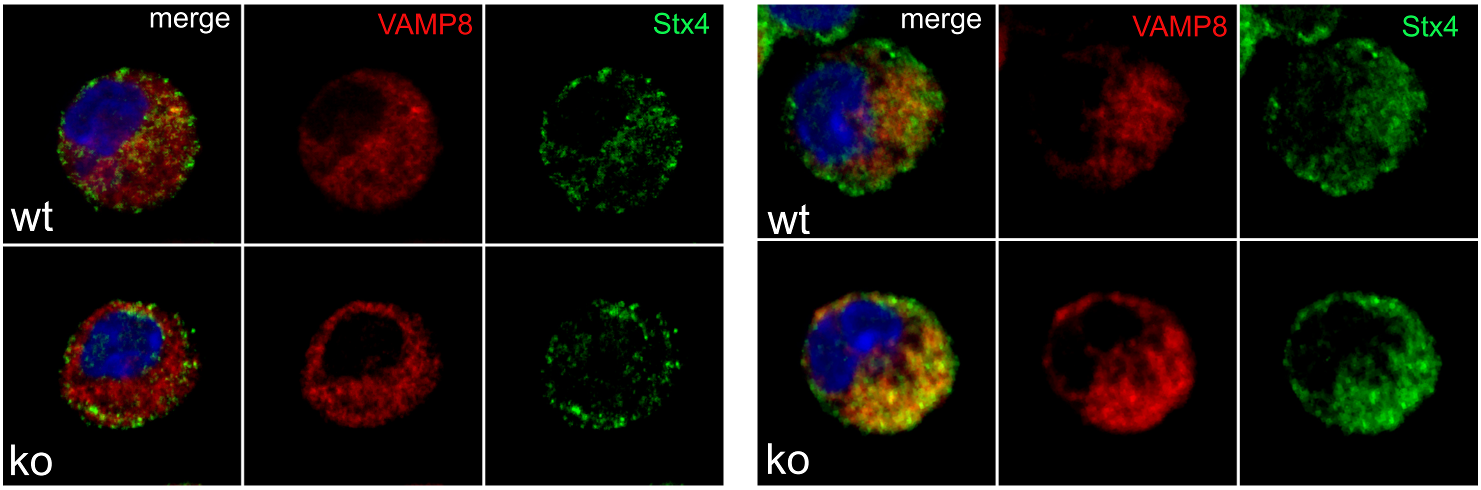

### Supplemental Figure 2

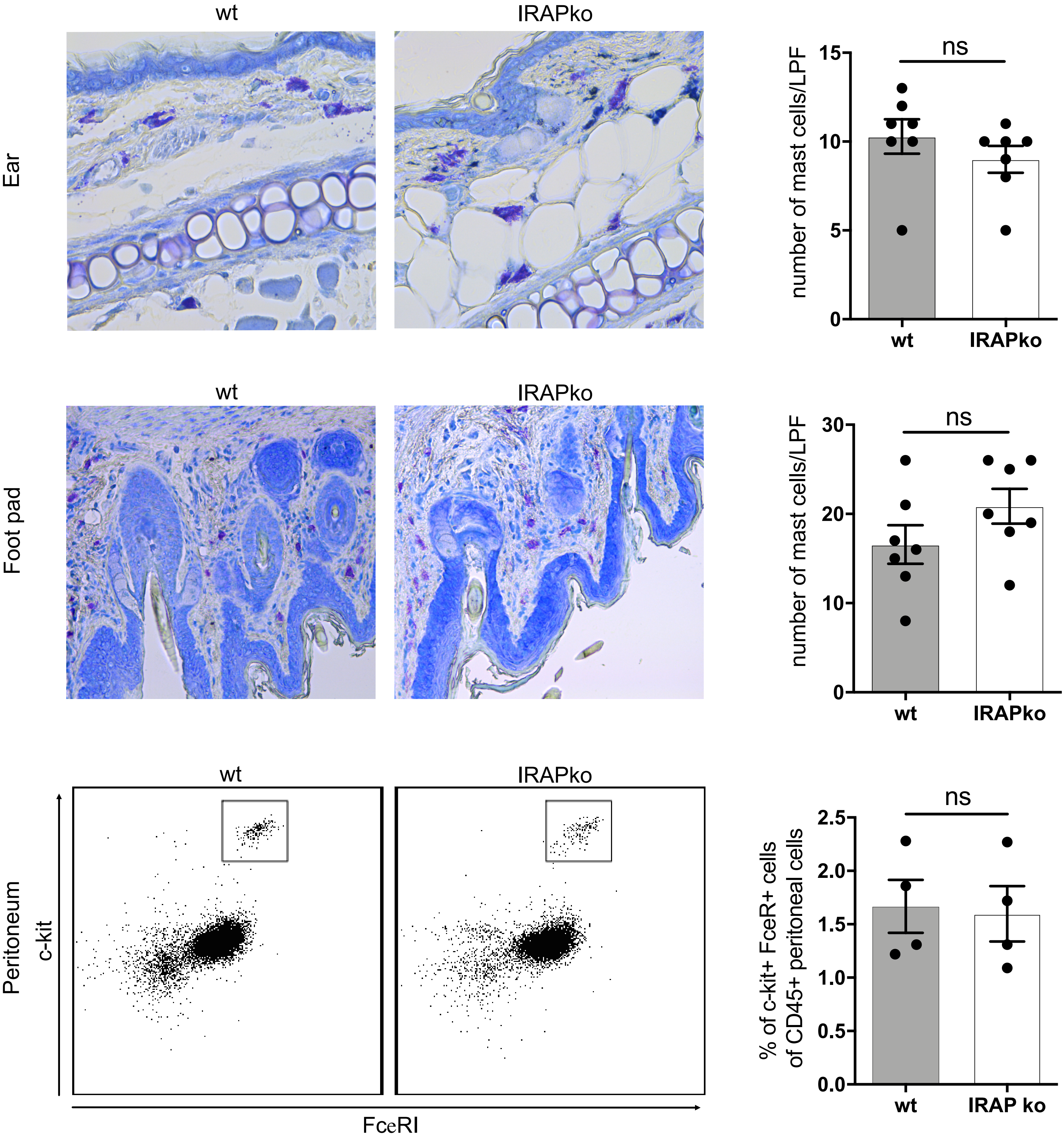

### Supplemental Figure 3

**Suppl. Fig.3**

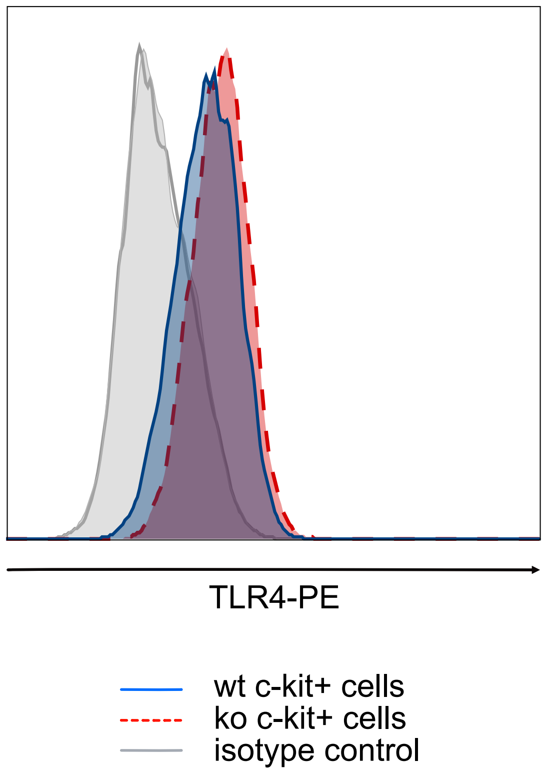

### Supplemental Figure 4

#### Suppl. Fig. 4

**A**

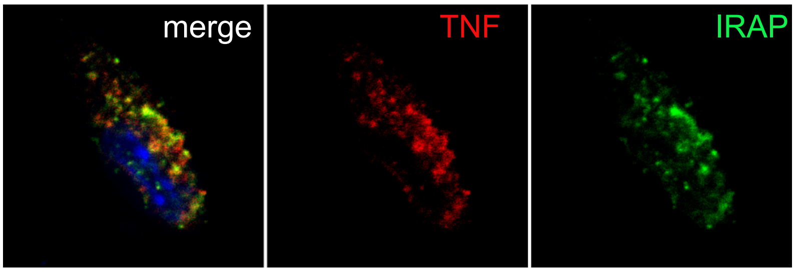


**B**

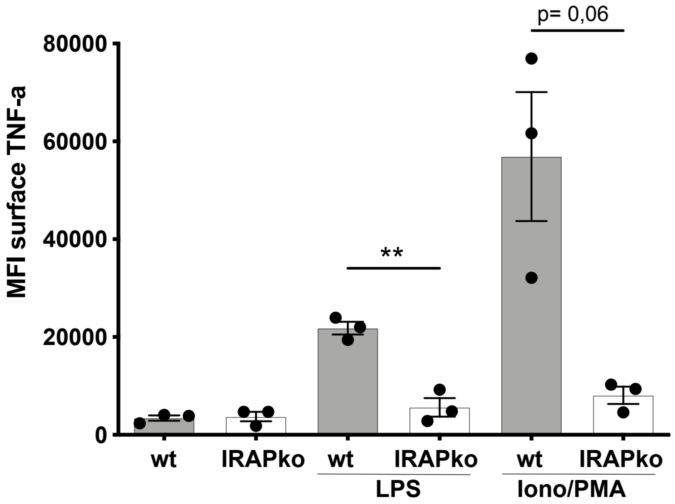

### Supplemental Figure 5

#### Suppl. Fig.5

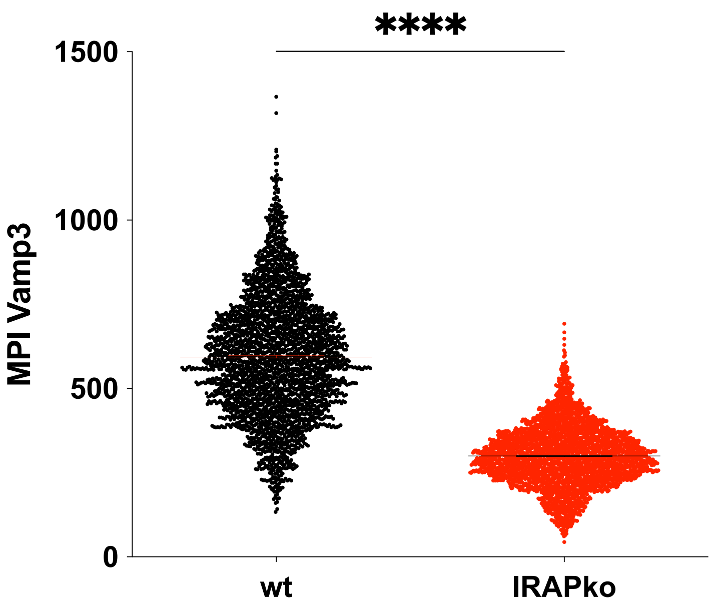

### Supplemental Figure 6

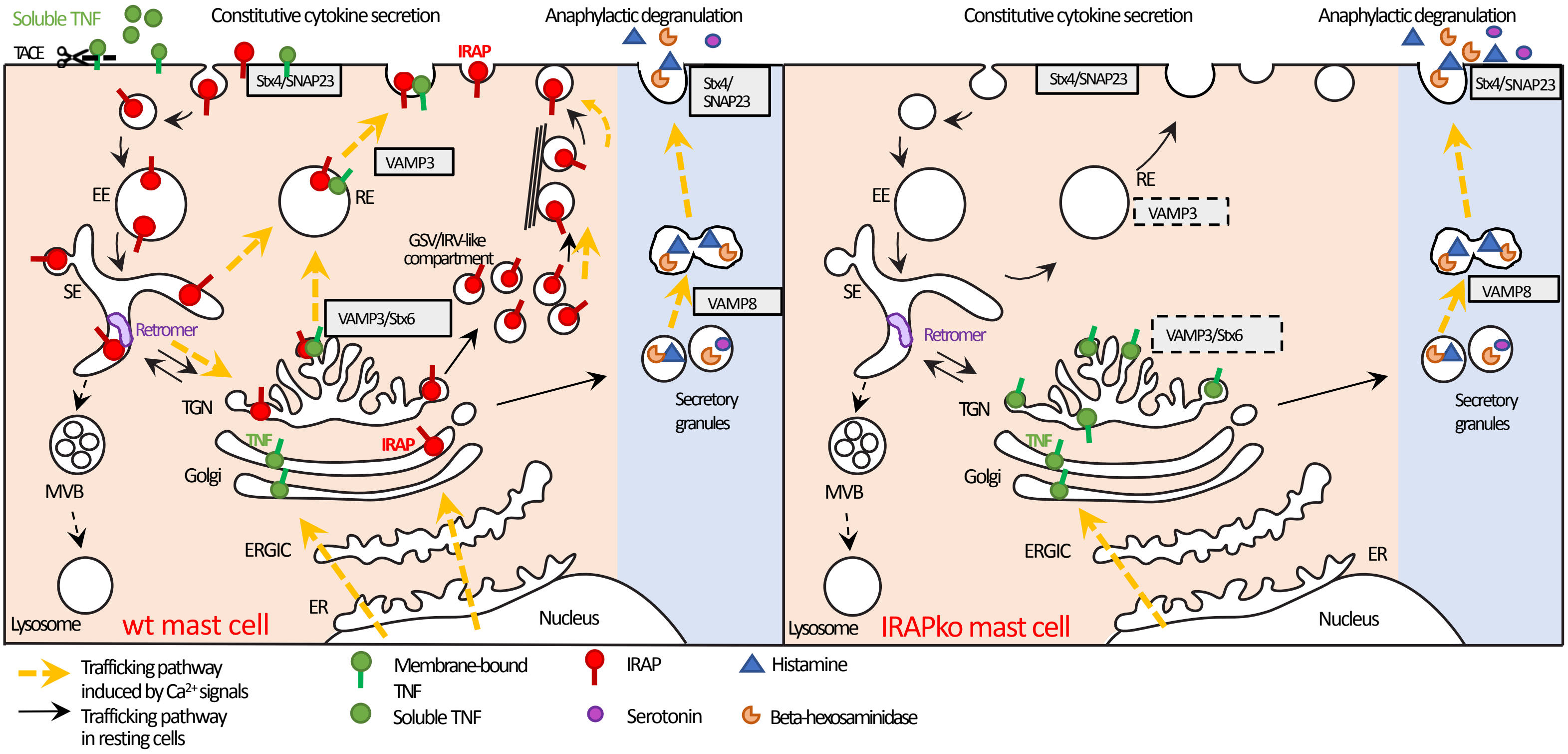

### Supplemental Figure 7

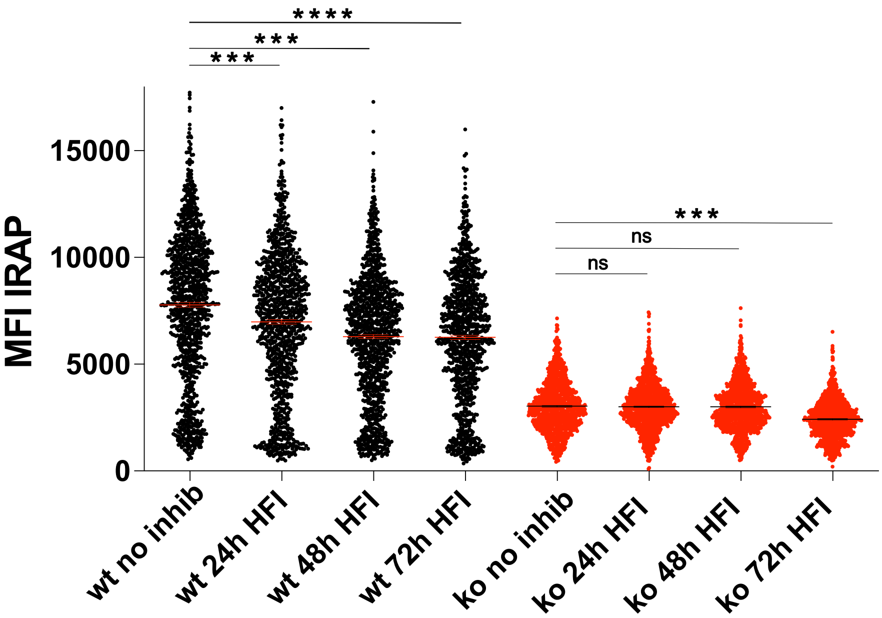
